# Genetic architecture of miRNA expression in human brain and its contribution to brain disorders

**DOI:** 10.64898/2026.01.15.699788

**Authors:** Arun H Patil, Anandita Rajpurohit, Yong Kyu Lee, Carly Montoya, Carrie Wright, Geo Pertia, Thomas M Hyde, Joel E Kleinman, Joo Heon Shin, Daniel R. Weinberger, Taeyoung Hwang

**Affiliations:** Lieber Institute for Brain Development, Baltimore, MD, USA; Department of Neurology, Johns Hopkins University School of Medicine, Baltimore, MD, USA; Department of Psychiatry and Behavioral Sciences, Johns Hopkins University School of Medicine, Baltimore, MD, USA; Department of Genetic Medicine, Johns Hopkins University School of Medicine, Baltimore, MD, USA; Solomon H. Snyder Department of Neuroscience, Johns Hopkins University School of Medicine, Baltimore, MD 21205, USA

## Abstract

MicroRNAs (miRNAs) regulate nearly all protein-coding genes and play critical roles in gene regulation, yet the mechanisms governing miRNA regulation remain poorly understood. Here, we examined the genetic architecture of miRNA expression in 995 human brain tissues spanning four regions and two ancestries (African and European) from neurotypical controls and individuals with three psychiatric disorders (schizophrenia, major depressive disorder, and bipolar disorder). We found that miRNA expression is highly dynamic across brain regions, with region-specific differences exceeding those attributable to ancestry or psychiatric diagnosis. Through miRNA expression quantitative trait loci (miR-eQTL) analyses, we identified genetic variants associated with miRNA expression in a region- and ancestry-dependent manner. Overall, miRNAs exhibited heritability levels comparable to protein-coding genes and were positively co-regulated with their host transcripts. Notably, miRNA-regulating genetic variants were enriched in enhancers active in oligodendrocytes and in binding sites of the transcription factor OLIG2, suggesting miRNA-mediated gene regulation in oligodendrocyte lineage. Finally, we identified 15 miRNAs as likely causative factors for psychiatric and neurodegenerative disorders such as major depressive disorder and Alzheimer’s disease. Together, our results reveal the genetic underpinnings of miRNA regulation in the human brain and suggest that miRNAs serve as key intermediaries linking genetic variation to complex neuropsychiatric and neurological phenotypes.

## Introduction

MicroRNA (miRNA) is a small non-coding RNA that represses gene activities in a post-transcriptional context. A single miRNA can target hundreds of mRNAs, exerting broad influence across complex gene regulatory networks^1,2^. In the nervous system, miRNAs play critical roles in neural development, synaptic plasticity, and neuronal activity^3,4^. Dysregulation of miRNA has been implicated in numerous brain disorders ^5–7^.

The biogenesis and molecular functions of miRNAs are well established^8,9^. Briefly, primary miRNA transcripts are sequentially processed by the nucleases Drosha and Dicer in the nucleus and cytoplasm, respectively, to generate about 22-nucleotide mature miRNAs. These mature miRNAs guide the RNA-induced silencing complex (RISC) to complementary target transcripts, promoting mRNA degradation or translational repression^8,9^. Despite this detailed understanding of miRNA function, the mechanisms that regulate miRNA expression themselves remain far less understood, particularly in human brain tissue where diverse and dynamic repertoires of miRNAs are involved in exceptionally complex gene regulation^10,11^. This complexity, combined with the cellular heterogeneity and limited accessibility of brain tissue^12^, has limited our understanding of how miRNA abundance is controlled genetically and epigenetically in the human brain.

Recent advances in genome technologies now allow the systematic investigation of genetic effects on RNA regulation by integrating genotype data with expression traits. Several studies have identified microRNA expression quantitative trait loci in human tissues, demonstrating that common genetic variants can influence miRNA expression ^13–22^. However, these studies have substantial limitations. First, relatively few have focused on brain tissues, and most available data are restricted to a single brain region such as the dorsolateral prefrontal cortex^13^ or are based on modest sample sizes (<100 samples)^14^, leaving unclear whether there are brain regional specificities in miRNA regulation. Second, previous analyses often examined miRNAs in isolation, without comparison to other gene classes such as protein-coding genes, which limits insight into the broader architecture of transcriptional regulation^13,14,18,20^. Third, nearly all studies have been conducted in individuals of European ancestry, despite evidence that regulatory variants and allele frequencies differ substantially across populations^23,24^. Collectively, these gaps have hindered a comprehensive understanding of the genetic regulation of miRNA expression in the human brain.

In this study, we systematically investigated the genetic architecture of miRNA expression in 995 human brain tissues across four brain regions and two ancestries (African and European), encompassing neurotypical controls and individuals diagnosed with major psychiatric disorders. We first characterized miRNA expression patterns across brain regions, ancestries, and three psychiatric conditions including schizophrenia, major depressive disorder, and bipolar disorder. We then performed miRNA expression quantitative trait loci (miR-eQTL) analyses, associating ∼10 million imputed SNPs with 566 highly expressed miRNAs in each region and ancestry group, and in parallel conducted gene expression QTL (eQTL) analyses using matched mRNA-seq data. By comparing the genetic architectures of miRNA and mRNA expression and assessing their co-regulation with host genes, we delineated shared and distinct patterns of regulatory control across brain regions and ancestries. Finally, we integrated miR-eQTL results with 165 genome-wide association studies (GWAS) to identify miRNAs potentially contributing to psychiatric and neurodegenerative disorders. Together, these analyses provide a comprehensive view of how genetic variation influences miRNA expression in the human brain and its potential impact on the molecular basis of neuropsychiatric and neurological diseases.

## Results

### miRNA expression landscapes across human brain region, ancestries, and psychiatric disorders

To characterize miRNA expression profiles in human brain tissue, we performed small RNA-seq on 995 postmortem brain tissues comprising 502 samples from 294 European (CAUC) and 493 samples from 201 African American (AA) individuals across four brain regions: dorsolateral prefrontal cortex (DLPFC), medial prefrontal cortex (mPFC), hippocampus, and caudate nucleus (Fig. 1A, Supplementary Table 1). These tissues were selected from the LIBD (Lieber Institute for Brain Development) brain repository to have matched genotype and mRNA data (Supplementary Fig. 1A). Sequencing reads were processed with miRge, our alignment software for small RNA-seq ^25^, with miRBase v22.1^26^ for annotation (see Methods). The number of retained miRNAs after QC was as follows (Supplementary Fig. 1B): DLPFC (1477), mPFC (1471), hippocampus (1465), and caudate (1449). A Uniform Manifold Approximation and Projection (UMAP)^27^ projection (Supplementary Fig. 1C) revealed distinct clustering of brain regions, with substantial overlap between DLPFC and mPFC, consistent with their shared neocortical identity.

**Figure 1.**
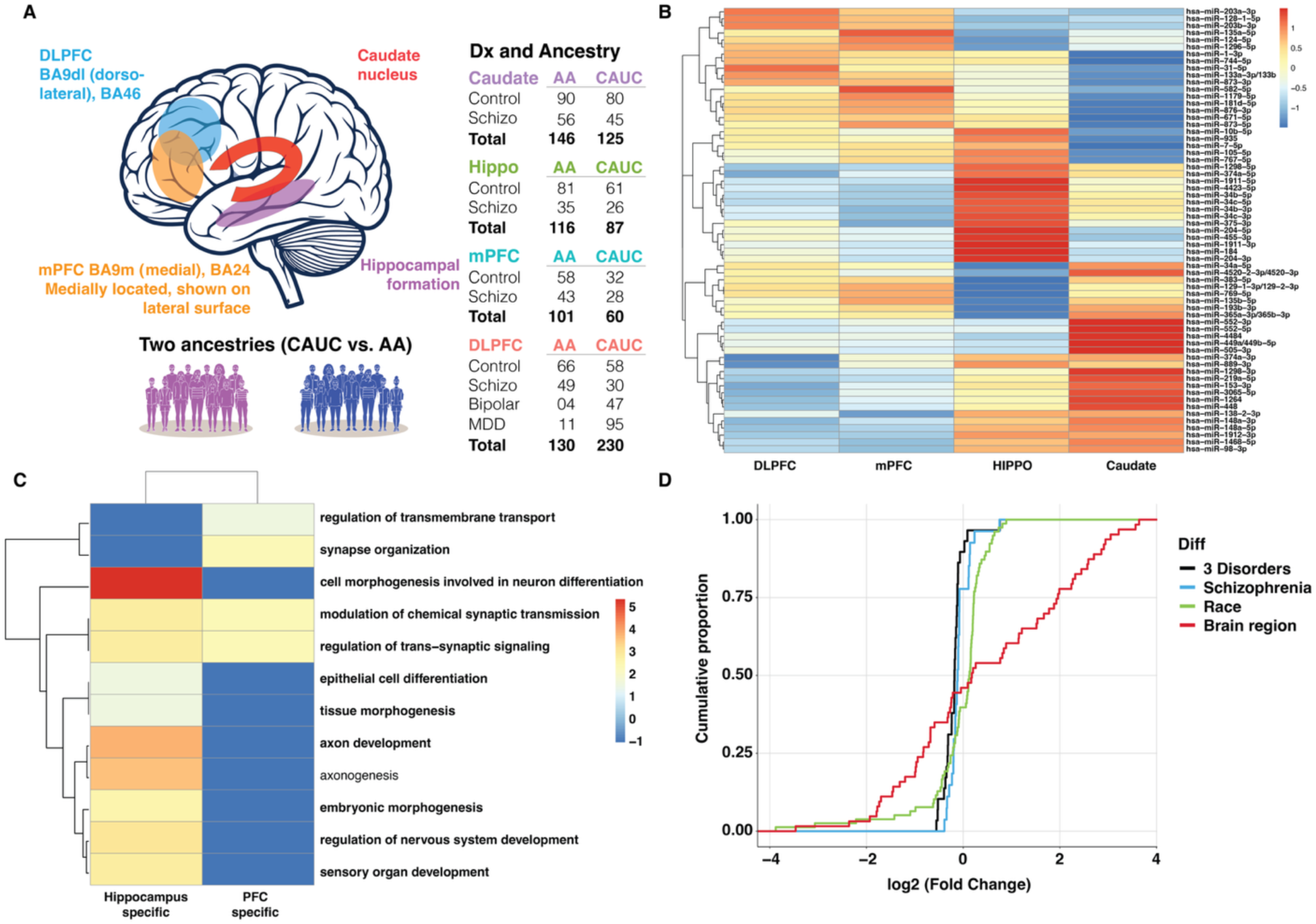
miRNA expression landscapes across brain regions, ancestries, and psychiatric disorders. (A) Overview of the 995 brain tissue samples analyzed in this study. (B) Heatmap of miRNAs differentially expressed across four brain regions. The color gradient represents scaled expression levels (z-score) in each row. (C) Significantly enriched Gene Ontology (GO) biological process terms for region-specific miRNA targets. The color gradient reflects –log_10_ (p-value). (D) Cumulative distributions of fold changes from differential expression analyses. Red: largest differences across DLPFC, mPFC, hippocampus, and caudate; Green: differences between European (CAUC) and African (AA) ancestries; Sky blue: control vs. schizophrenia; Black: largest differences among control, schizophrenia, bipolar disorder, and major depressive disorder (MDD).

To identify miRNAs differentially expressed across prefrontal cortex (DLPFC and mPFC), hippocampus and caudate, we used a likelihood ratio test (LRT) implemented in DESeq2^28^, incorporating RNA integrity number (RIN), sequencing depth, sex, ancestry, and primary diagnosis as covariates of a linear model. This analysis identified 789 miRNAs that are significantly expressed differentially across these four brain regions (FDR < 0.01, Supplementary Table 2). Of these, 63 miRNAs exhibited strong regional differences, defined by a maximum mean expression difference > 0.5 between the highest- and lowest-expressing regions (Fig. 1B). We further identified region-specific miRNAs based on the pairwise comparison between the selected regions. Specifically, for each miRNA, we compared the brain region with the highest mean expression with the region exhibiting the next-highest mean expression level and quantified the difference with Cohen’s d. We defined miRNAs showing higher Cohen’s d (>1) as region-specific miRNAs (Supplementary Figure 2): the prefrontal cortex exhibited three highly specific miRNAs, while the hippocampus and caudate showed seventeen and three, respectively.

To explore the biological roles of region-specific miRNAs, we performed Gene Ontology (GO)^29^ enrichment analysis on their predicted target genes (target score > 80 in mirDB^30^). Twelve biological processes were significantly enriched (q < 0.05; gene ratio > 0.05), primarily associated with neuronal features (Fig. 1C). Region-specific patterns were also observed: prefrontal cortex–enriched miRNAs targeted genes involved in synapse organization, modulation of chemical synaptic transmission, and trans-synaptic signaling while hippocampal miRNAs were enriched for nervous system development and neuron differentiation. We didn’t detect any significant terms for caudate-specific miRNAs.

We next examined whether miRNA expression varied by ancestry or psychiatric diagnosis. Using a Wald test in DESeq2^28^, we identified 78 miRNAs differentially expressed between ancestries. Among these, miR-577-5p showed the strongest difference, with ∼43–100% higher expression in individuals of European ancestry compared to African Americans (Supplementary Fig. 3A). Conversely, miR-1299 was consistently relatively under-expressed in European samples across all regions (Supplementary Fig. 3B). These miRNAs showing the significant differences between CAUC and AA showed a similar trend when expression differences were quantified selectively in AA samples by the regression of the expression levels on the estimated proportion of AA ancestry within each brain of the AA donors, suggesting strong genetic effects on the expression differences in ancestry (Supplementary Fig. 3C).

For psychiatric disorders, we applied LRTs across major depressive disorder (MDD), bipolar disorder (BP), schizophrenia (SCZ), and controls in the DLPFC, and Wald tests for SCZ vs. control across the four brain regions. Note that bipolar and depression cases were available only in DLPFC. These analyses identified 29 and 27 differentially expressed miRNAs (FDR < 0.01), respectively. For example, miR-212-5p was consistently downregulated in SCZ across three brain regions (Supplementary Fig. 4A) while miR-451a was significantly upregulated in SCZ across three brain regions (Supplementary Fig. 4B). However, overall, diagnosis-related differences were smaller in magnitude than regional or ancestry-based differences in miRNA expression (Fig. 1D).

### Genetic regulation of miRNA expression

We next investigated the genetic regulation of miRNA expression by performing miRNA expression quantitative trait loci (miR-QTL) analyses ^31^, linking miRNA expression to genetic variants located within ±0.5 Mb of each miRNA transcription start site (TSS). Analyses were performed separately for each of the four brain regions and two ancestry groups. Because some miRNAs share identical mature sequences and can originate from multiple genomic loci, potentially confounding QTL assignment, the analysis was restricted to uniquely annotated miRNAs. Across brain regions, we found 849 and 923 (LD-independent) loci that harbor significant miR-QTLs (FDR<0.05) in CAUC and AA, respectively. The numbers of emiRs (miRNAs with at least one significant cis-QTL) ranged from 35 in the European hippocampus to 224 in the European DLPFC (Fig. 2A, Supplementary Table 3). The DLPFC exhibited the largest number of emiRs, while the caudate had the fewest, consistent with variation in sample size across regions (Supplementary Fig. 5A). When statistical significance was evaluated by permutation test, the similar pattern was also observed with smaller number of detected emiRs (Supplementary Fig. 5B). Several eMiRs, including miR-211-5p and miR-3615-3p, were consistently detected across all brain regions and ancestries (Supplementary Fig. 6 and Supplementary Table 3).

**Figure 2.**
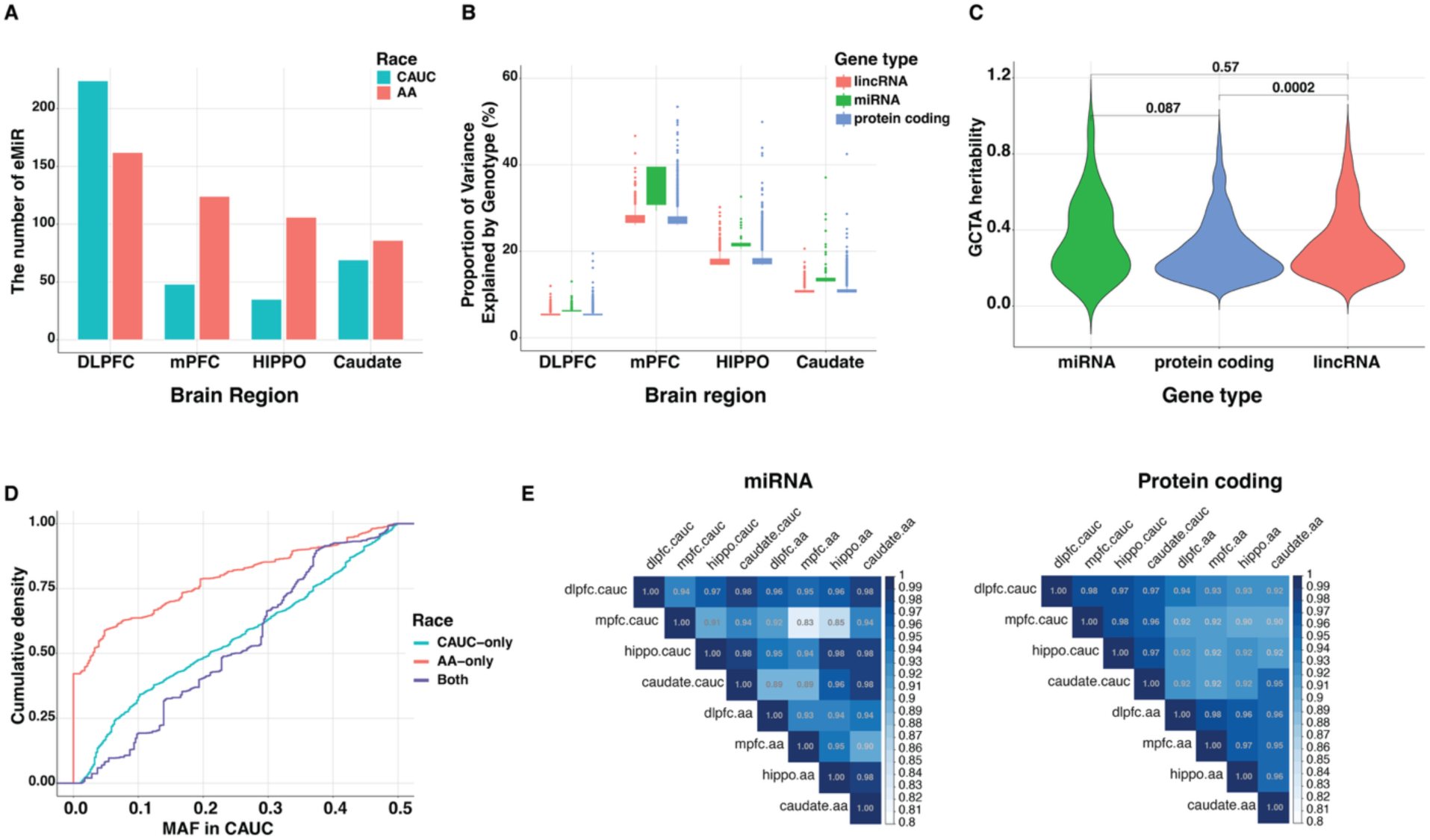
Genetic regulation of miRNA expression. (A) Number of emiRs (miRNAs with significant cis-miR-eQTL variants) identified across brain regions and ancestries. (B) Distribution of the proportion of variance explained (PVE) by lead variants for each expressed miRNA. (C) cis-heritability of miRNA expression estimated using GCTA. (D) Cumulative proportions of minor allele frequencies (MAF) in European ancestries for the significant cis-miR-eQTL variants found only in European ancestries (CAUC, blue), African ancestries (AA, red) and both (Both, purple). (E) Degree of sharing of significant lead miR-eQTL signals among pairs of brain regions and ancestries. The concordance between brain regions relative to ancestries is less clear with miRNA, compared with protein coding genes.

For the most significant cis-variants of every eMiRs, the percent variance explained (PVE) ranged from 6.1% to 46.3% (mean = 14.0%) in European DLPFC samples (Fig. 2B) although PVE estimates can be inflated by smaller sample sizes in some brain regions. Compared to protein-coding or long noncoding (lncRNA) genes, miRNAs exhibited, on average, higher PVEs across brain regions and ancestries, suggesting that miRNA expression varies more predictably with genotype than mRNA expression (Fig. 2B and Supplementary Fig. 7A). To validate this observation, we used GCTA^32^ to estimate SNP-based cis-heritability by aggregating the variance explained by all SNPs within ±1 Mb of each miRNA. Across brain regions and ancestries, miRNAs showed heritability levels comparable to or higher than protein-coding genes (Fig. 2C and Supplementary Fig. 7B), indicating that common genetic variants substantially influence miRNA expression in the brain.

We next assessed the tissue and ancestry specificity of miR-QTL effects. First, we observed 150 miRNA eQTLs (emiRs) in DLPFC that were detected exclusively in individuals of AA ancestry, despite the smaller AA sample size compared with European ancestry (N = 130 vs. 230, respectively). Examination of allele frequencies revealed that, for the majority of these associations, the minor alleles at the lead eQTL variants (variants with highest significance) were present in AA but absent or extremely rare in CAUC (Fig. 2D). These findings indicate that differences in allele frequency across ancestries substantially contribute to the observed ancestry-specific miRNA eQTL signals, highlighting the importance of diverse populations for uncovering genetic regulation of miRNA expression. Second, we used Multivariate Adaptive Shrinkage (MASH) analysis^33^, which leverages correlation structures between tissues to estimate shared or tissue-specific effects. Briefly, among the effects that are significant in at least one of the comparing tissues, MASH computed what fraction of those have an estimated (posterior mean) effect size within a factor of one another (here 0, effectively considering only sign). MASH revealed strong concordance (larger than 0.9) of miR-QTL effects across brain regions, similar to the patterns observed for mRNA eQTLs (Fig. 2E). However, despite this high overall concordance, miRNA eQTLs exhibited a lower degree of cross-tissue sharing than protein-coding gene eQTLs in both European and African ancestries, indicating a more tissue-specific regulatory architecture for miRNAs (Fig. 2E).

### Characterization of miR-eQTLs and their regulatory context

To gain insight into the functional mechanisms underlying miRNA regulation, we annotated cis-miR-QTL variants with known genomic and regulatory features. We first intersected the lead variants of every miRNAs with chromatin contexts of human brain tissues obtained from NIH Roadmap epigenetics data^34,35^ (Fig. 3A). The emiR variants were significantly enriched in promoter and enhancer regions, consistent with previous findings^13,36^. Interestingly, these variants tended to reside farther from the transcription start site (TSS) than eQTLs for protein-coding genes, for example, on average 396 kb vs. 347 kb in the European DLPFC cohort (Supplementary Fig. 8). This result suggests that distal enhancer elements may exert a larger influence on miRNA expression than promoter-proximal variants.

**Figure 3.**
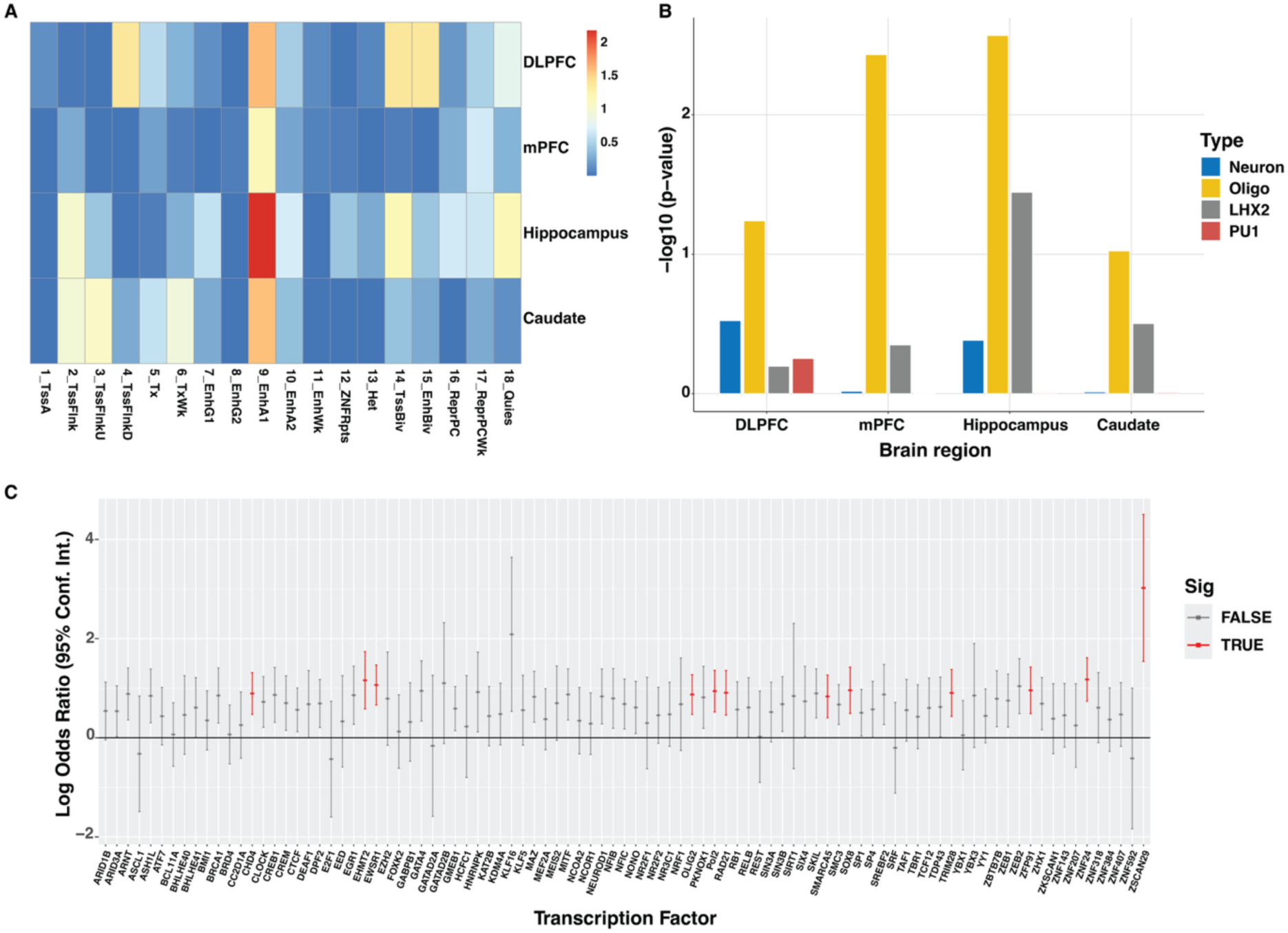
Regulatory context of miR-eQTL enrichment. (A) Enrichment of CAUC miR-eQTLs across 18 chromatin states determined by chromHMM in human brain tissues. The color gradient reflects –log_10_(p-value). (B) Enrichment of CAUC miR-eQTLs within cell type–specific enhancers from major brain cell types. (C) Enrichment of CAUC miR-eQTLs in transcription factor (TF) binding regions. Significantly enriched TFs are indicated in red.

We next asked the cell-type specificity of miR-QTLs using enhancer annotations from major brain cell types^37^. Among neurons, astrocytes, oligodendrocytes, and microglia, miR-QTLs showed strongest enrichment in oligodendrocyte-specific enhancers (Fig. 3B). To identify potential transcriptional regulators mediating this effect, we examined overlap between miR-QTLs and transcription factor (TF) binding sites. Notably, OLIG2, a master regulator of oligodendrocyte differentiation and myelination, emerged as one of top significantly enriched TFs (Fig. 3C). These results suggest that miRNA expression in the brain is at least partly regulated through oligodendrocyte-associated transcriptional programs, highlighting a potential role for miRNA-mediated mechanisms in myelination and glial function.

Because many miRNAs are embedded within host genes, we next examined the extent of genetic co-regulation between intragenic miRNAs and their host transcripts. Among the 394 emiRs, 339 (86%) were located within annotated protein-coding genes that were detected in our samples. To test whether these intragenic miRNAs share regulatory variants with their host genes, we performed colocalization analysis using coloc^38^, a Bayesian method that estimates the posterior probability of shared causal variants. We identified 16 miRNA–host gene pairs with evidence of colocalization (posterior probability H₄ > 0.5) across 4 brain regions. These pairs exhibited significantly higher positive expression correlations between miRNAs and their host mRNAs than non-colocalized pairs (Supplementary Fig. 9). However, host genes were not predicted targets of their embedded miRNAs, suggesting that the observed co-expression reflects shared transcriptional regulation rather than miRNA-mediated feedback repression.

### Evaluating miRNA’s association with brain disorders

To investigate the potential contribution of miRNA expression to complex brain traits, we performed transcriptome-wide association studies (TWAS)^39^ and Mendelian Randomization (SMR) ^40^ using 87 GWAS summary datasets^41^ from individuals of European ancestry, spanning psychiatric, neurological, and behavioral phenotypes (Supplementary Table 4). For stringent discovery, we integrated multiple TWAS approaches: we first used FUSION to predict miRNA–trait associations based on cis-eQTL effects on miRNA expression, and then applied SMR and HEIDI tests to identify associations consistent with shared causal variants rather than linkage. Across all analyses, we identified 15 miRNAs significantly associated with a range of brain disorders, including bipolar disorder, schizophrenia, and Alzheimer’s disease (Fig. 4A and Supplementary Table 5). The predicted targets of these miRNAs are involved in neuron differentiation and axon development (Supplementary Fig. 10). We also performed TWAS for AA ancestry using 56 GWAS datasets selected from a recent study of the VA-MVP cohort (Supplementary Table 6)^42^. We found 4 miRNAs predicted to cause motor movement-related traits (Supplementary Table 5).

**Figure 4.**
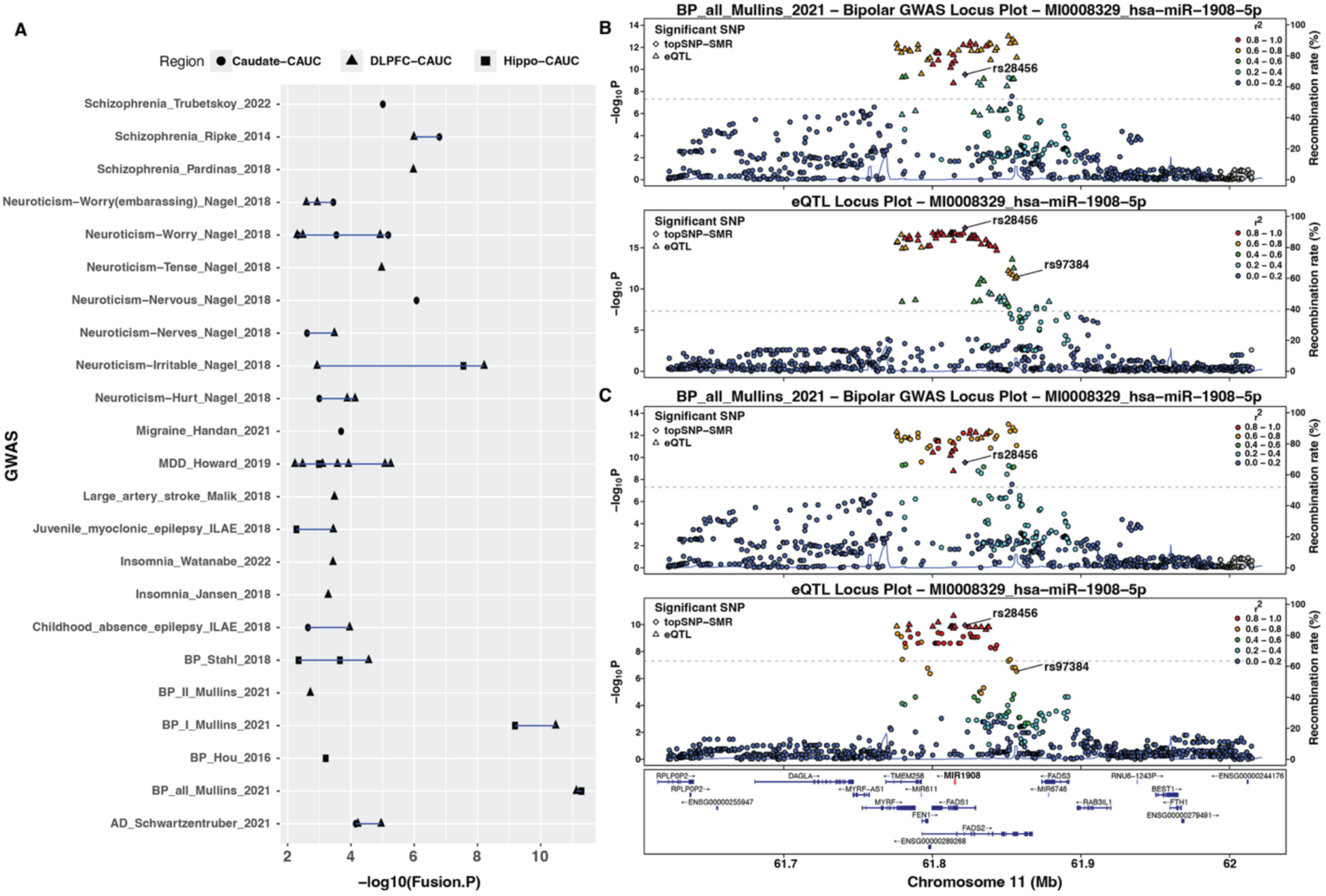
Transcriptome-wide association study (TWAS) of miRNA–trait associations. (A) Traits showing significant TWAS associations. Only the top miRNA–trait association for each brain region is displayed. (B) TWAS result for miR-1908-5p in the dorsolateral prefrontal cortex (DLPFC). The upper panel shows the corresponding GWAS association signals, and the lower panel shows the miR-eQTL signals. (C) Same as (B), but for the hippocampus.

Among 15 miRNAs associated as causative factors in brain disorders in European ancestry, the strongest signal was observed between miR-1908-5p and bipolar disorder, extending a previously reported association in DLPFC ^13^ to both DLPFC and hippocampus (Fig. 4B). For Alzheimer’s disease, miR-200a/b was predicted to decrease Alzheimer’s disease risk, providing genetic support for a protective effect of this miRNA that has been variably reported in experimental studies^43^. For schizophrenia, we identified miR-2682-5p, which resides within the MIR137HG locus, one of the most strongly implicated genomic regions in schizophrenia GWAS^44,45^ (Supplementary Fig. 11). While miR-137 did not show significant cis-heritability of expression in our data (GCTA p-value of miR-137-3p: 0.5) and was therefore excluded from the TWAS analysis, our TWAS analysis predicted that increased expression of miR-2682-5p was positively associated with schizophrenia risk. This finding is consistent with the lead eQTL SNP showing negative effects on both expression and GWAS association. Although miR-2682-5p is expressed at substantially lower levels than miR-137 (Supplementary Fig. 12), its significant association suggests that multiple miRNAs within the MIR137HG locus may contribute to schizophrenia risk.

## Discussion

In this study, we generated a genome-wide dataset comprising genetic variants, mRNA expression and miRNA expression to investigate the potential impact of miRNA and the genetic regulation of miRNAs in human brain tissues. This resource allows a systematic analysis of miRNA expression across four brain regions spanning two ancestries and multiple psychiatric disorders. We show that variability of miRNA expression that is explained by common genetic variations is comparable to or slightly greater than that of protein-coding genes. As miRNAs regulate broad numbers of target genes affecting many cellular processes, they might be expected to be robust to non-genetic factors, potentially increasing the contribution of genetic components especially for a subset of miRNAs. However, we also observed weaker cross-tissue concordance of genetic regulation for miRNAs compared with protein-coding genes, indicating a more tissue-specific regulatory architecture. Although ancestry-specific regulatory effects were detected, many of these differences were attributable to ancestry-specific allele frequency distributions rather than distinct regulatory mechanisms.

miRNAs have been reported to exhibit strong spatiotemporal regulation across development. Consistent with this, we observed brain region-specific miRNA expression patterns. Several miRNAs are preferentially expressed in specific brain regions, with distinct functional enrichments. Notably, hippocampus-specific miRNAs preferentially targeted genes involved in neuronal differentiation and nervous system development, whereas miRNAs enriched in DLPFC or mPFC were more strongly associated with synaptic pathways. Recent studies suggest that adult neurogenesis in the human hippocampus is limited compared with that of rodents, raising questions about its molecular regulation in humans. Interestingly, one of the hippocampus-specific miRNAs identified in our analysis, miR-184, has been previously shown to repress adult neurogenesis in mice^46^. The hypothesis that miR-184 may contribute to the regulation of adult hippocampal neurogenesis in humans warrants further functional investigation.

Many miRNAs are located within host genes, expected to be sharing their transcriptional regulation with host genes. Consistent with this, we observed positive correlations of expression between intragenic miRNA and host mRNAs when they are impacted by co-localized genetic variations. But the colocalized host mRNAs do not tend to be predicted targets of intragenic miRNAs although a previous study identified two examples of host genes as targets of intragenic miRNAs ^13^. These findings do not support feed-forward regulatory mechanism for target genes through miRNAs, at least in adult brain tissues.

Many psychiatric disorders have a strong genetic component, but most studies so far attribute the heritability to protein-coding genes, resulting in the limited accountability. By integrating eQTL and GWAS statistics, we identified many miRNAs causally associated with psychiatric disorders. Among them, we noted miR-1908 which is strongly associated with bipolar and major depressive disorders. Our multi region analysis has replicated this association not only in DLPFC as reported previously ^13^ but also in hippocampus, increasing the level of replication. We also focused on a causal association between miR-2682 and schizophrenia. miR-2682 resides within the MIR137HG locus, one of the most strongly associated schizophrenia risk loci identified by GWAS. This locus harbors both miR-2682 and miR-137, which might be expected to be co-regulated as they are embedded within the same host gene, MIR137HG^47^. miR-137 has been a primary focus of functional studies due to its relatively higher expression levels. For example, previous experiments on miR-137 overexpression is associated with schizophrenia-related molecular perturbations in mouse and in cell models^48–51^. However, our results demonstrate that miR-2682 expression, despite its low expression overall, is highly correlated with miR-137, and increased miR-2682 expression is causally associated with schizophrenia risk, while miR-137 was not. This apparent discrepancy may be explained by the fact that miR-137 showed no significant cis-heritability of expression in our data and was therefore excluded from the TWAS analysis. Consistent with our findings, a recent study reported that miR-2682 was significantly upregulated in schizophrenia patients compared to controls and had greater magnitude of changes than miR-137^52^. These findings suggest that two miRNAs may independently or jointly contribute to schizophrenia susceptibility, but requires experimental studies to disentangle their individual and combined functional roles.

While most brain tissue studies of psychiatric disorders have focused on neurons, accumulating evidence have shown that glial cells are also involved in disease pathogenesis. In particular, oligodendrocytes, which are responsible for elaborating myelin sheaths around axons to strengthen the signal communications between neurons, have emerged as a cell type associated with psychiatric risk^53,54^ and is affected by stressful experiences ^55^. For example, Liu et al. reported that protracted social isolation decreases myelin gene products and nuclear heterochromatin formation, thus inducing transcriptional and ultrastructural changes in OLs of the PFC, which ultimately led to impaired adult myelin formation^56^. Also, oligodendrocyte maturation was reported to be dependent on social experience ^57^. Consistent with these studies, we found that genetic loci associated with miRNA expression are enriched in enhancers active in oligodendrocytes. Also, the genetic variants are enriched with the binding regions of a transcription factor, OLIG2 that is critical for oligodendrocyte differentiation and maturation. It will be interesting to see miRNA’s functional roles in oligodendrocyte and their dysregulation in psychiatric disorders.

While our analysis focused on the genetic regulation of miRNA expression across brain regions and ancestries, our dataset also includes miRNA expression profiles from individuals with several psychiatric disorders. Although we observed only modest effect sizes between control and disorder groups at the level of individual miRNAs, future studies could extend these findings by examining other brain regions, miRNA–gene network interactions and co-regulatory modules to reveal broader regulatory mechanisms underlying psychiatric disease. Overall, we provide a comprehensive resource of miRNA expression profiles from human brain tissues that will facilitate future investigations into molecular etiology of psychiatric or other brain disorders.

## Method

### Brain tissues

Tissues were obtained from dorsolateral prefrontal cortex (DLPFC) of individuals of African and European ancestries diagnosed with schizophrenia, bipolar disorder, major depressive disorder (MDD), and neurotypical controls. In addition, tissues from caudate, hippocampus, and medial prefrontal cortex (mPFC) were obtained from individuals of African and European ancestries diagnosed with schizophrenia and from neurotypical controls. The procurement of samples and analysis involving human brain regions were subject to following all the ethical requirements and informed consents obtained from the next of kin as described previously^58^. The detailed information of samples is provided in Supplementary Table 1.

### Genotype imputation and quality control

Initital genotype calling was performed for 680 individuals from LIBD Brain repository using several Illumina SNP BeadChip microarrays (112 samples with HumanMap650Y, 435 with Human1M-Duo, 91 with HumanOmni5-Quad, and 42 with Infinium Omni2.5-8). Genotype phasing and imputation was performed on separate batches for each of the different SNP arrays, using the TOPMed ^59^ with the hg38 reference panel. We finally filtered the merged imputed genotype data to only keep variants having MAF > 0.01 and converted to BED format using Plink v1.9 ^60^, generating the variant calling file (VCF ^61^) for 12,339,188 SNPs. The quality control for individual genotype data was performed using PLINK 2.0 as previously described ^36^, briefly, the SNPs were filtered for Hardy-Weinberg equilibrium (--hwe 1e-6), minor allele frequency (--maf 0.01), individual missing genotype rate (--mind 0.1), and variant missing genotype rate (--geno 0.05). For this project, analyses were restricted to the adult subset (n = 495 of 680 individuals).

### Small RNA-seq

For each sample, 500ng total RNA was used as input for the PerkinElmer NEXTFLEX Small RNA-seq Kit v3, followed by our previous study^62^. Libraries were prepared according to the manufacturer’s protocol, following the procedure for gel-free size selection. For the PCR step of the protocol, 17 cycles were used for all samples, based on the manufacturer’s recommendations and previous studies^62^. Fifty-base-pair single-end sequencing was performed with the Illumina HiSeq 3000.

### Preprocessing of small RNA-seq and miRNA quantification

Illumina 4N Unique molecular Identifiers (UMI) were extracted from sequencing reads and used to remove PCR duplicates after the 3-prime Illumina adapters were removed using Cutadapt^63^ (v 4.9). The processed small-RNA reads were aligned and assigned by miRge ^25^. Briefly, sequencing reads were aligned to the custom-built RNA-species library annotations provided with miRge using Bowtie^64^ (v 1.3.1) and human miRBase^26^ database (release 22.1) was used as reference miRNA.

We initially generated 1448 small RNA-seq datasets of CAUC and AA brain tissues. We filtered low quality samples based on the proportion of miRNA-assigned reads (miRcountRate>=0.1), the number of detected miRNAs (miRnum<500) and the availability of RNA-seq. We also filtered low-expressed or sparsely-detected miRNAs whose max counts are smaller than 2 or whose counts are found less than 10 tissues, generating 1481 miRNA counts for every tissue. After selecting adult tissues for this project, we further filtered samples that are not clustered into 3 major brain regions (DLPFC/mPFC, Hippocampus, Caudate) based on UMAP (version 0.2.10.0)^27^ with the normalized counts obtained from DESeq2’s variance stabilizing transformation (VST)^28^. Outlier samples identified through these analyses were manually inspected and excluded from downstream analyses (n = 60 individuals), resulting in 995 tissues.

### Differential expression

Differential expression (DE) analysis of miRNAs was performed across brain regions, diagnostic groups, and ancestries using DESeq2^28^. Specifically, likelihood ratio tests (LRT) were applied to evaluate expression differences attributable to brain region, while controlling for relevant covariates. The full model included RIN, sequencing depth, sex, ancestry, primary diagnosis, and brain region, whereas the reduced model excluded the brain region term. Quantitative covariates were scaled, and categorical variables were encoded as factors. For differential expression analysis between ancestries (Caucasian and African American), pairwise Wald tests were conducted. When using a continuous estimate of African American (AA) genetic ancestry as an independent variable, we obtained ancestry proportion values from a previously published study with overlapping donors^58^. Briefly, the African ancestry admixture proportion for each individual was estimated using the STRUCTURE program based on ancestry-informative SNP markers. For diagnosis, Wald tests was used to assess differences between Schizophrenia and Controls across the Hippocampus, Caudate, and mPFC regions, while the LRT was applied in the DLPFC to evaluate differential expression across multiple diagnostic groups, including Bipolar Disorder, Major Depressive Disorder (MDD), Schizophrenia, and Controls.

### MicroRNA expression quantitative trait loci (miR-QTLs)

The miRQTL analysis was performed across four brain regions and two ancestries, Dorsolateral prefrontal cortex (DLPFC) with 230 Caucasians (CAUC) and 130 African American (AA) individuals, Caudate (n= 125 CAUC; 146 AA individuals), Hippocampus (n=87 CAUC, 116 AA individuals) and Medial prefrontal cortex (mPFC, n= 60 CAUC and 101 AA individuals). Abundantly expressed miRNAs whose read counts were greater than or equal to 10 in at least 10 samples were retained for analysis. Then, read counts were normalized using DESeq2’s variance stabilizing transformation (VST)^28^. Residuals were then generated from the VST-transformed data by fitting a linear model that adjusted for scaled quantitative covariates, including RIN, sequencing depth, and age.

Cis-miRNA QTL mapping was performed with the residuals using TensorQTL^31^ (v1.0.10) in both *permutation* and *nominal* modes, with a cis-window of 1 Mb. Analyses were conducted separately within each ancestry-specific brain region. In each analysis, the covariates included (i) Genotype-derived principal components (PCs) - top 10 PCs computed using PLINK2.0^60^ after linkage disequilibrium (LD) pruning (window = 50 kb, step = 5, *r*² < 0.2) to account for population structure; (ii) Expression-derived PCs - top 10 PCs obtained using the prcomp function in R (version 4.3.3) (center = TRUE, scale = TRUE) from normalized miRNA residuals, representing latent technical and biological variation (miRNAs with zero variance across samples were excluded prior to PCA); (iii) Categorical covariates - sex and primary diagnosis, encoded as a model matrix.

### Heritability Estimation

SNP-based heritability of miRNA expression was estimated using the Genome-wide Complex Trait Analysis (GCTA) tool^32^ (v1.94.1). Covariates from the eQTL analysis were processed to create appropriate quantitative and categorical inputs for GCTA. For each miRNA, a genetic relationship matrix (GRM) was constructed using SNPs within a ±1 Mb window around the gene locus (via gcta64 --make-grm), based on genotype data generated using PLINK2^60^. Heritability estimation was then performed using the GCTA-GREML approach to quantify the proportion of phenotypic variance attributable to common SNPs. To compare the genetic contribution of miRNAs with other RNA types, the GCTA-based heritability analysis was similarly extended to mRNA using the same methodological framework. The variance in expression explained by a single SNP was calculated under an additive model (2*f*(1-f)*b^2^/s_y_^2^,f: minor allele frequency, b: SNP effect size, s_y_^2^: trait variance in population).

### Multivariate adaptive shrinkage analysis (MASH)

MASH^33^ was used to identify shared and region-specific miR-QTL effects. Strong signals were obtained from the variants whose miR-QTL z-statistics were largest in 8 miR-QTL analyses (4 brain regions and two ancestries) while 50,000 randomly selected miR-QTLs were used to estimate the null correlation structure. Covariance matrices were derived via principal component analysis (PCA) and FLASH decomposition, complemented by canonical patterns representing shared and region-specific effects. MASH models were fit to the random data to learn mixture weights, then applied to strong signals to compute posterior summaries, including effect sizes and local false sign rates (LFSR).

### Colocalization between miRNA and host gene

We defined miRNAs as intragenic if they were located within protein-coding genes’ gene bodies and were on the same strand as a host gene, and as intergenic otherwise. We used coloc^38^ (version 5.2.3) to determine colocalization.

### Transcriptome-wide association study (TWAS)

TWAS^39^ pipeline for miRNA was implemented to link genetically regulated miRNA expression to complex traits. First, cis-heritability of miRNA expression was estimated using the Genome-wide Complex Trait Analysis (GCTA) tool ^32^. Heritable miRNAs were then integrated with genome-wide association study (GWAS) summary statistics using the FUSION TWAS framework ^39^ to identify putative miRNA–trait associations. Finally, causal relationships were evaluated using Summary-based Mendelian Randomization (SMR) ^40^, leveraging shared genetic architecture between miRNA expression and trait variation. The details are as follows: (1) LD Reference: The 1000 genome project reference data, generated by the New York Genome Center (NYGC)^65^, were used. European sample identifiers were matched to the LD reference panel provided by the TWAS-FUSION package. For African populations, family IDs corresponding to the following population codes were selected: African Caribbeans in Barbados (ACB), Americans of African Ancestry in SW USA (ASW), Esan in Nigeria (ESN), Gambian in Western Divisions in the Gambia (GWD), Luhya in Webuye, Kenya (LWK), Mende in Sierra Leone (MSL), and Yoruba in Ibadan, Nigeria (YRI); (2) GWAS summary data: GWAS summary statistics for the selected traits, including neurological, psychiatric, and neurodevelopmental disorders, were obtained from the Brain Catalog^41^. (3) FUSION: First, FUSION.compute_weights.R^39^ script was used to generate expression weights for each miRNA exhibiting significant SNP-based heritability, as determined by GCTA-GREML, across multiple brain regions and ancestral populations (see Heritability estimation). The FUSION script was executed using default settings, except for the --hsq_set parameter, which bypassed internal heritability estimation and instead used miRNA-specific heritability estimates derived from the GCTA analysis. Additionally, a covariate file consistent with that used in GCTA was provided via the --covar flag. Multiple testing correction was performed using the Benjamini-Hochberg method. An association was considered significant if it met the following criteria: Fusion p-value < 0.05, SNP-based heritability estimate (HSQ) > 0.01, eQTL model R² > 0.05, an absolute TWAS Z-score > 2 and FDR<0.05.

(5) Summary-based Mendelian Randomization (SMR): To further validate TWAS results, we employed SMR^40^ analysis. Results were considered significant if the SMR p-value was < 0.05 and the HEIDI test p-value was > 0.05, indicating a shared causal variant rather than linkage.

### Enrichment analysis

Gene ontology: Predicted miRNA target genes were obtained from the mirDB^30^, using a target score threshold >80 to define high-confidence targets. Gene Ontology (GO) enrichment analysis of biological processes was conducted using the enrichGO() function from the clusterProfiler R package^29^. Significance was determined at a *q*-value < 0.05, with *P*-values adjusted for multiple testing using the Benjamini–Hochberg procedure.

Genomic features: GARFIELD^66^ was used for the enrichment analysis of the selected variants for chromatin states calculated by CSREP^35^ with NIH Roadmap epigenetics brain profiles^34^, brain enhancer regions^67^ and transcription factor binding sites of DLPFC^68^. The significance threshold was determined within GARFIELD, which estimates the effective number of independent tests and applies a Bonferroni correction at the 5% level internally.

## Author Contributions

T.H., J.H.S. and D.R.W. conceptualized and supervised the research. T.M.H and J.E.K provided tissue resources, A.R. and J.H.S. generated the raw sequencing data with assistance of Y.K.L., C.M., and C.W.. G.P. processed genotype data. T.H. and A.H.P. carried out the computational analyses. T.H., A.H.P., A.R., and D.R.W. wrote the manuscript with the input from all authors.

## Acknowledgments

We thank the LIBD Neuropathology Section for their work in assembling and curating the clinical and demographic information and organizing the Human Brain Tissue Repository of the LIBD. We thank the families that have donated this tissue to advance our understanding of psychiatric disorders. The African Ancestry Neuroscience Research Initiative is a collaboration between the LIBD, Morgan State University, Duke University and members of the community led by Rev. Dr. A. C. Hathaway. We thank Dr. Siyu Pan for providing the curated GWAS results from the Brain Catalog database.

---

**Supplementary Figure 1.**
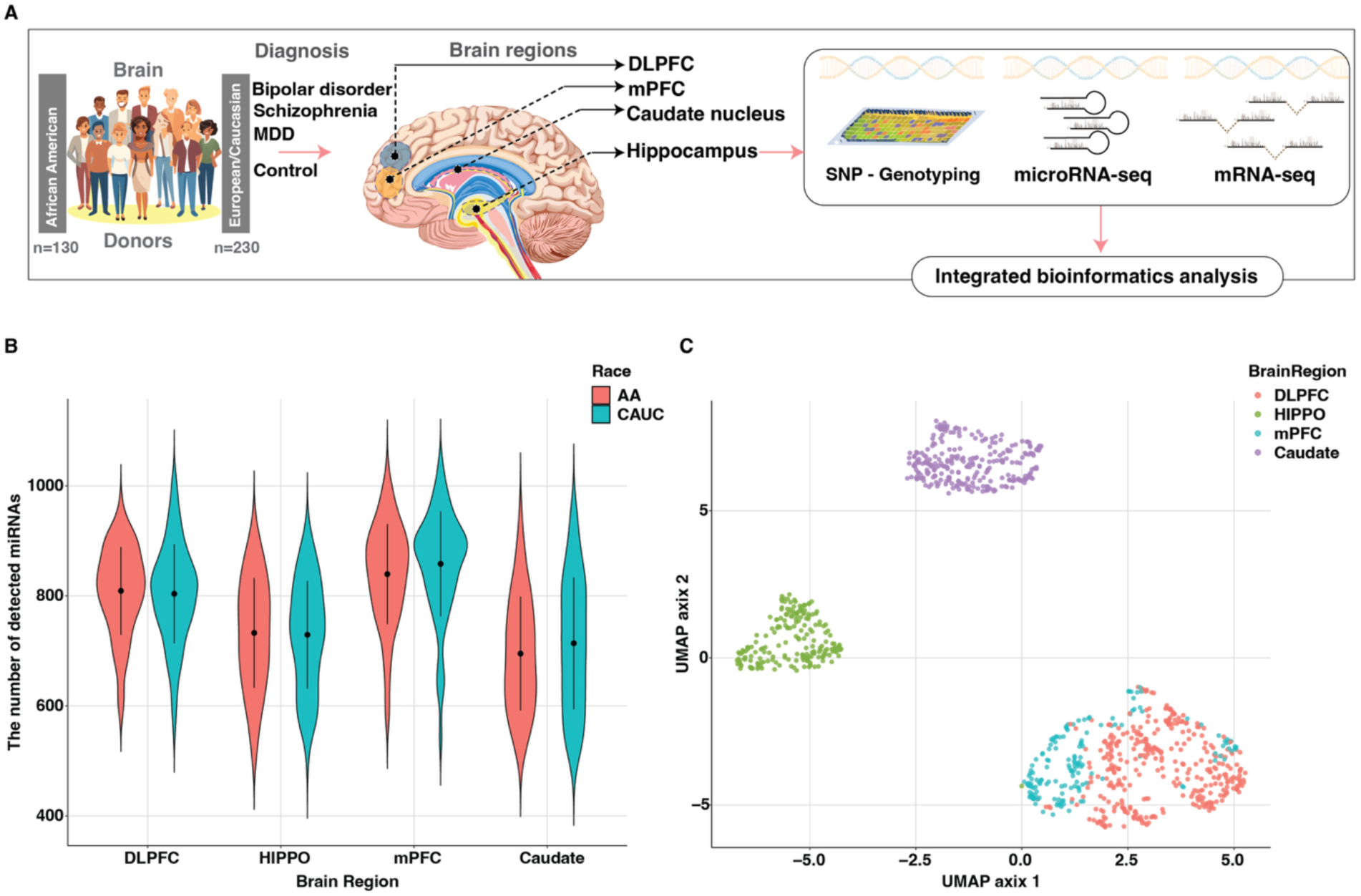
Overview of miRNA expression data and analysis. (A) Schematic overview of the study design, datasets, and analysis workflow. (B) Distribution of detected miRNAs (Reads Per Million>1) across ancestries and brain regions. (C) UMAP visualization of miRNA expression profiles showing clustering by brain region.

**Supplementary Figure 2.**
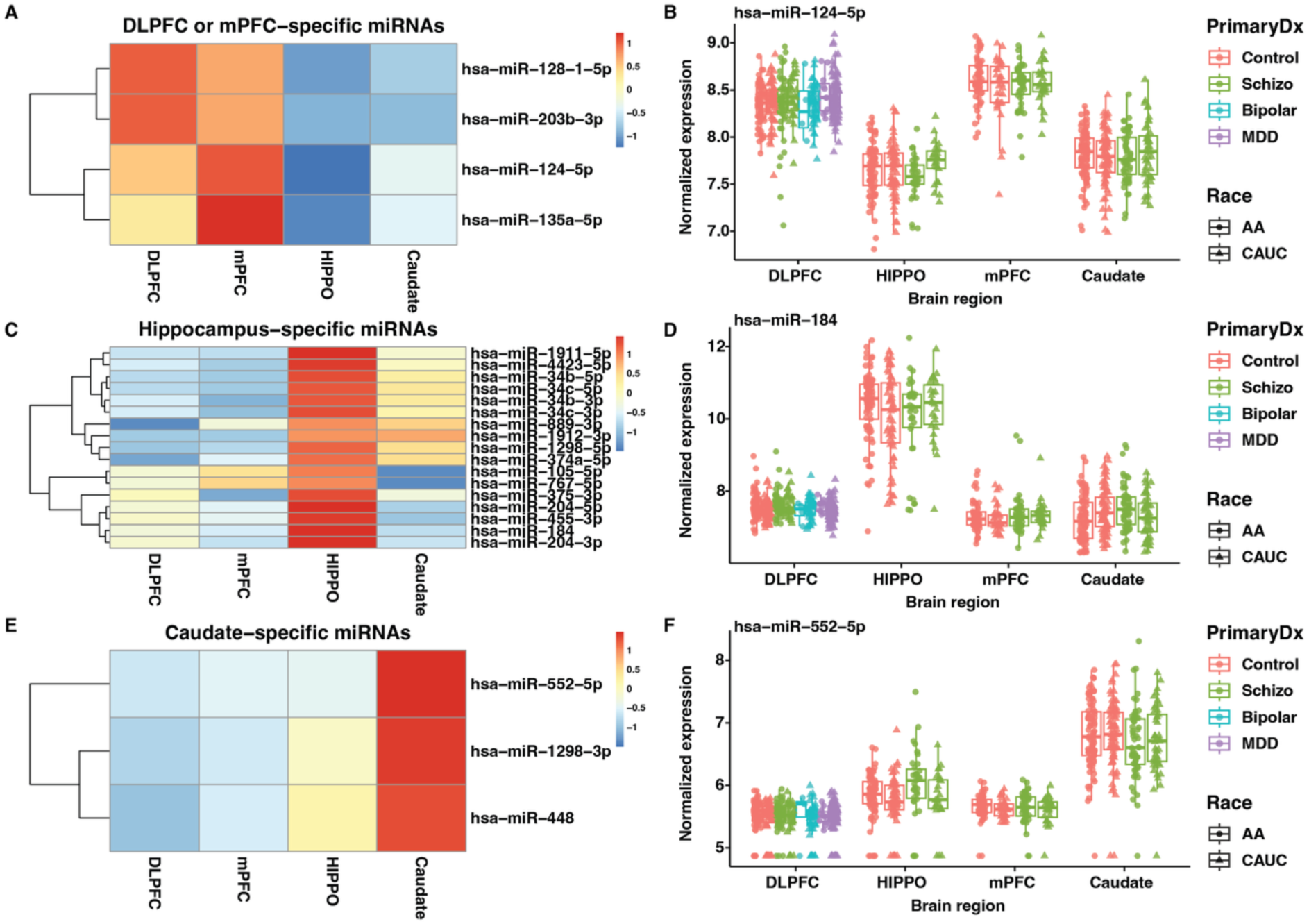
Brain region–specific miRNAs. miRNA expression of (A) prefrontal cortex tissues (DLPFC and mPFC), (C) hippocampus, and (E) caudate. The color gradient represents scaled expression levels (z-score) in each row. (B,D,F) Representative examples of region-specific miRNAs from each region, respectively.

**Supplementary Figure 3.**
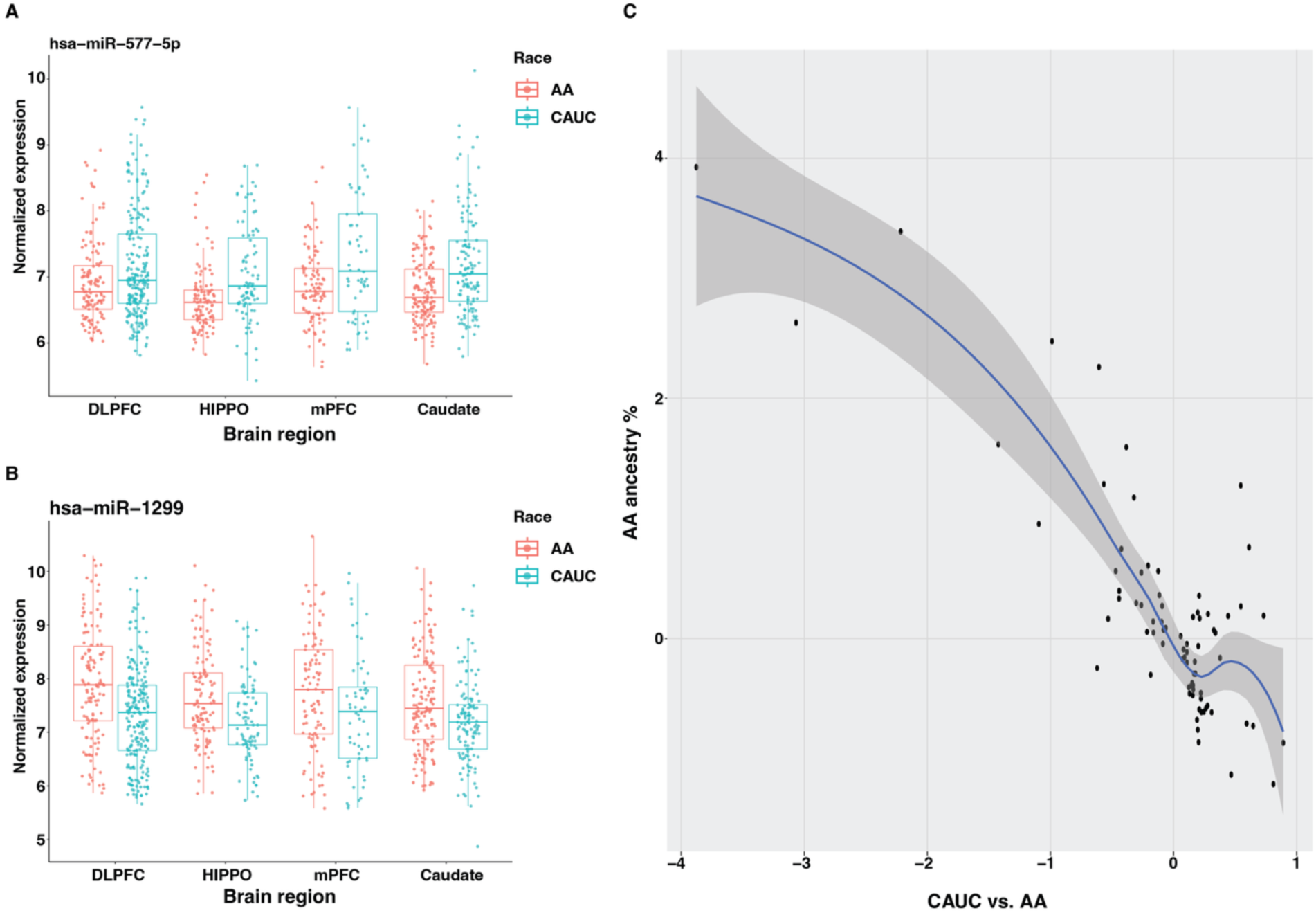
Differentially expressed miRNAs between European and AA ancestries. (A, B) Examples miRNAs showing differential expression between African American (AA) and Caucasian (CAUC) individuals. (C) The correlation between log2 fold changes of miRNA expressions between CAUC and AA ancestries: x-axis is log2 fold change estimated from the comparison between CAUC and AA ancestries, y-axis is log2 fold change estimated from the regression of miRNA expressions in AA on the admixture proportion of AA. Note that each dot is a miRNA.

**Supplementary Figure 4.**
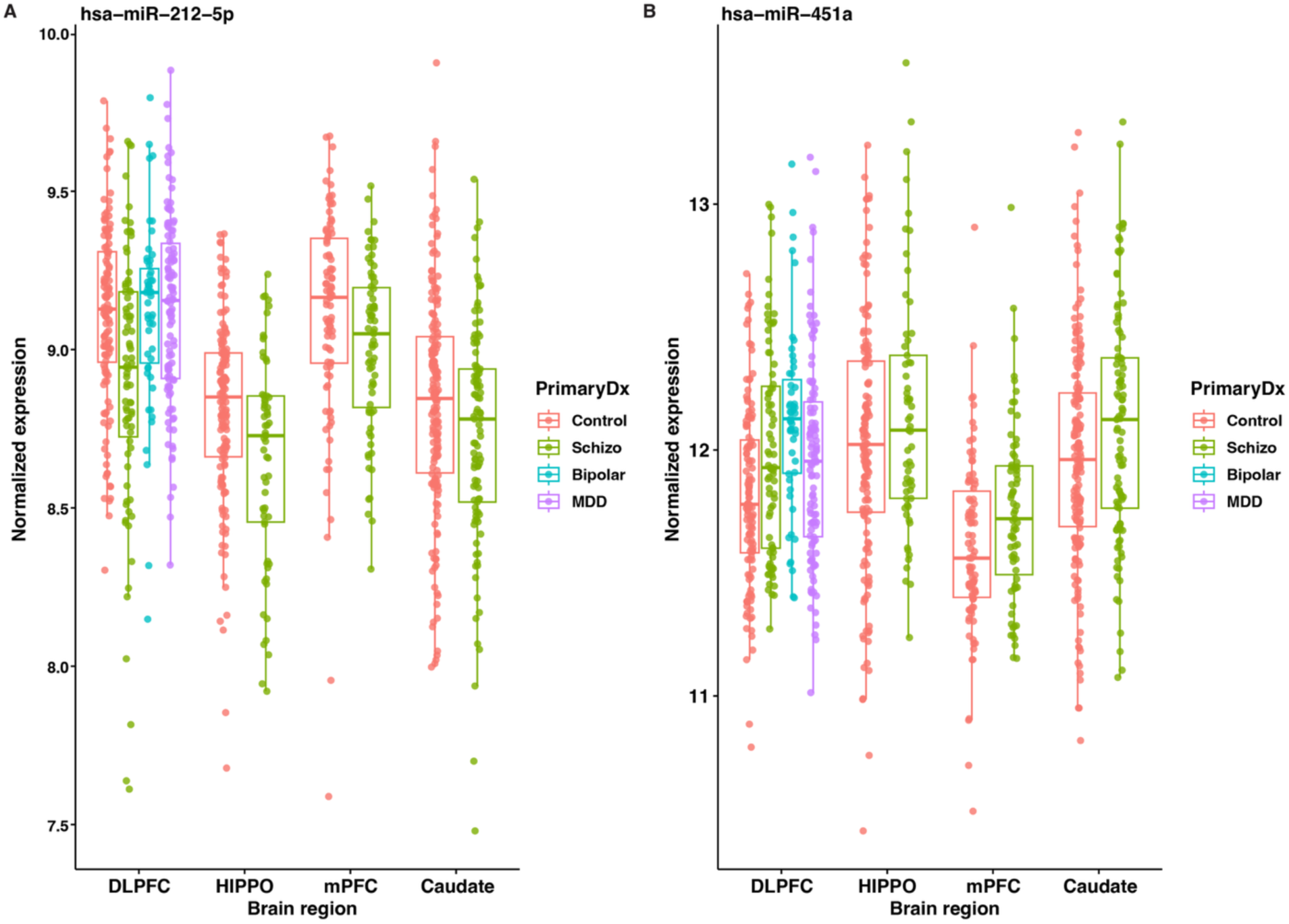
Differentially expressed miRNAs between control and schizophrenia. (A, B) Examples of miRNAs showing differential expression between control and schizophrenia individuals.

**Supplementary Figure 5.**
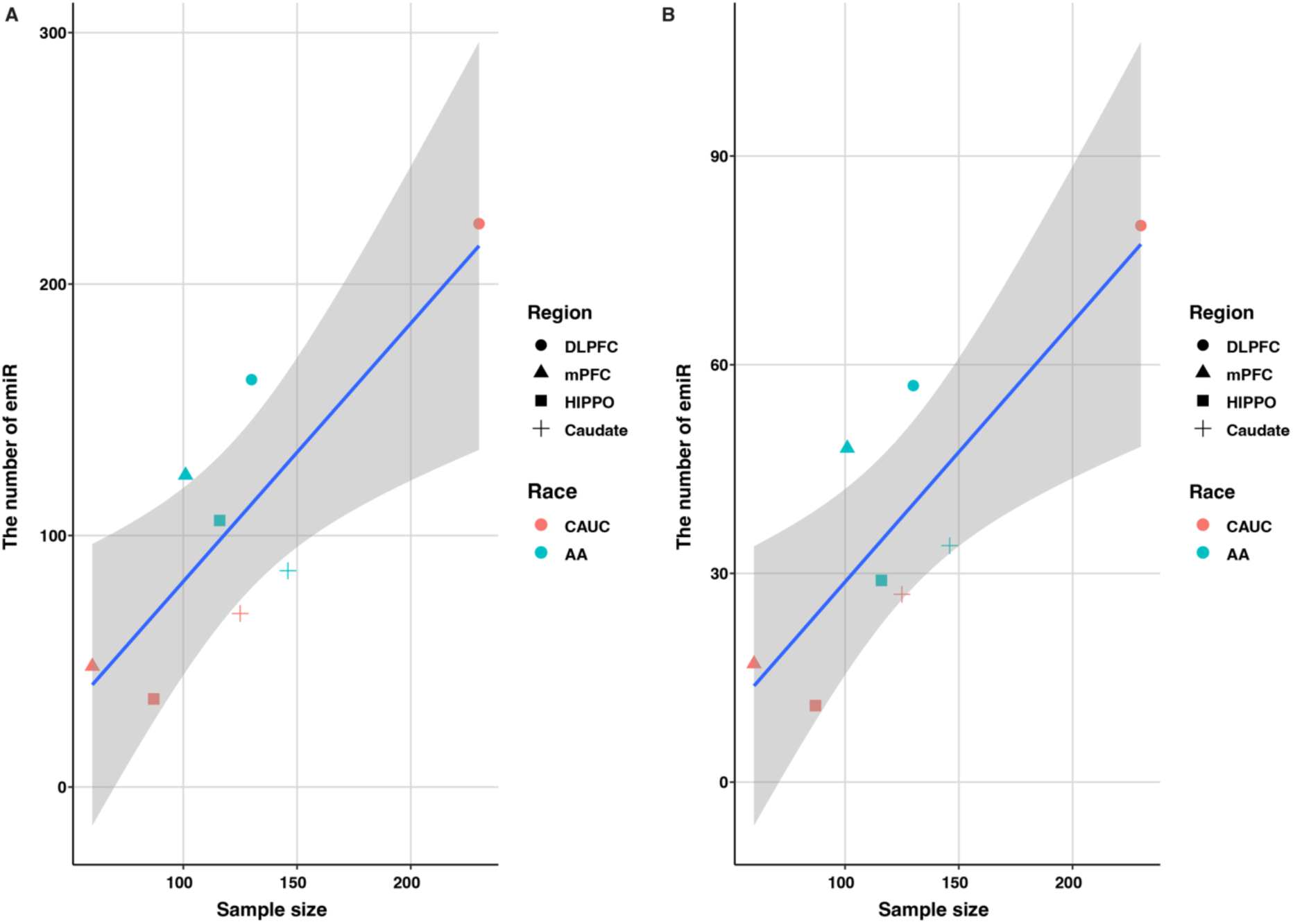
Correlation between sample size and the number of emiR. (A) The number of emiRs based on nominal p-value, (B) The number of emiRs based on p-value of permutation test.

**Supplementary Figure 6.**
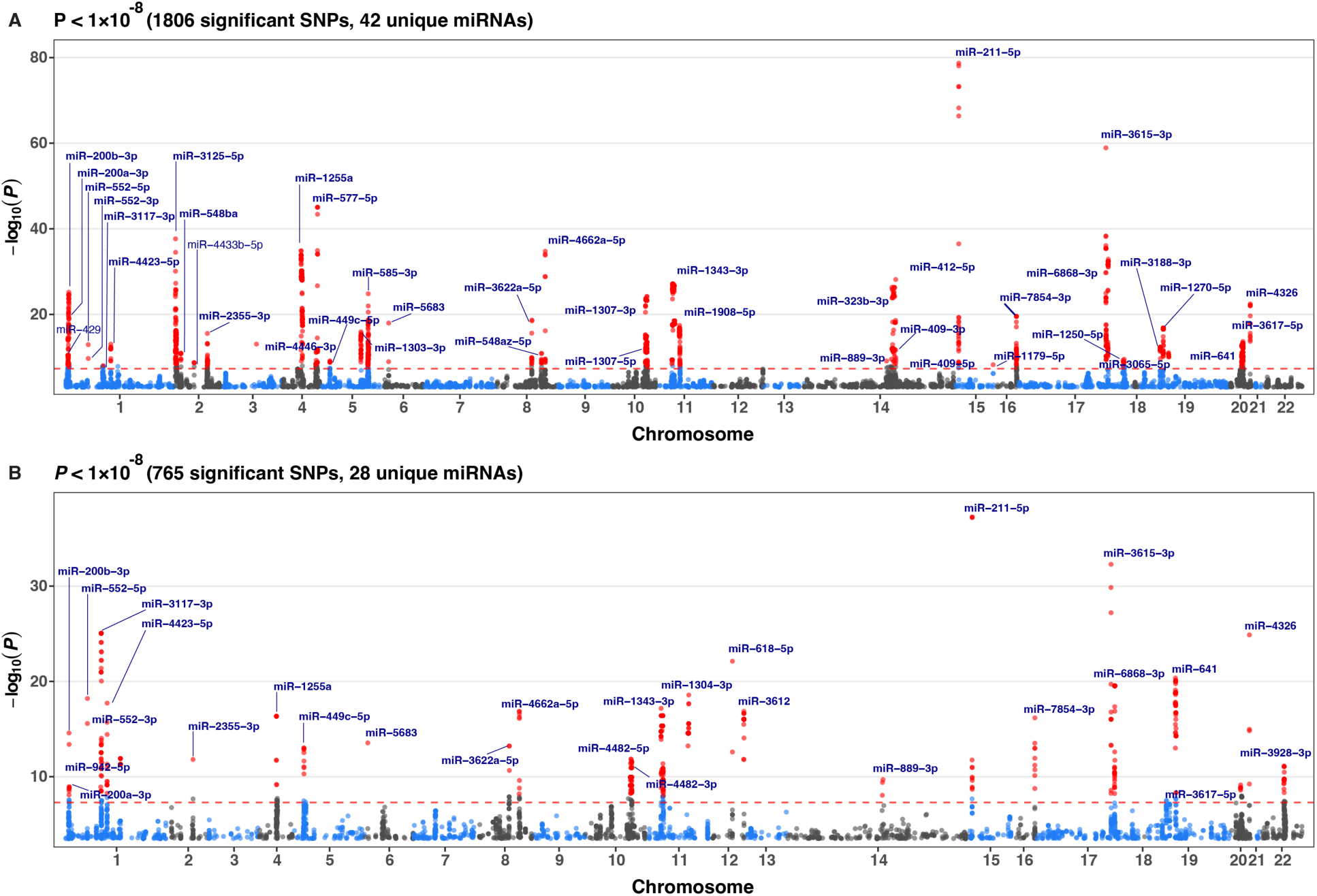
Manhattan plots of miR-eQTLs. (A) Results from the European (CAUC) cohort, combined across four brain regions. The y-axis represents −log₁₀ nominal P-values for variants with MAF > 0.2. For each locus, only the most significant SNP across regions is shown. (B) Same as (A), but for the African American (AA) cohort.

**Supplementary Figure 7.**
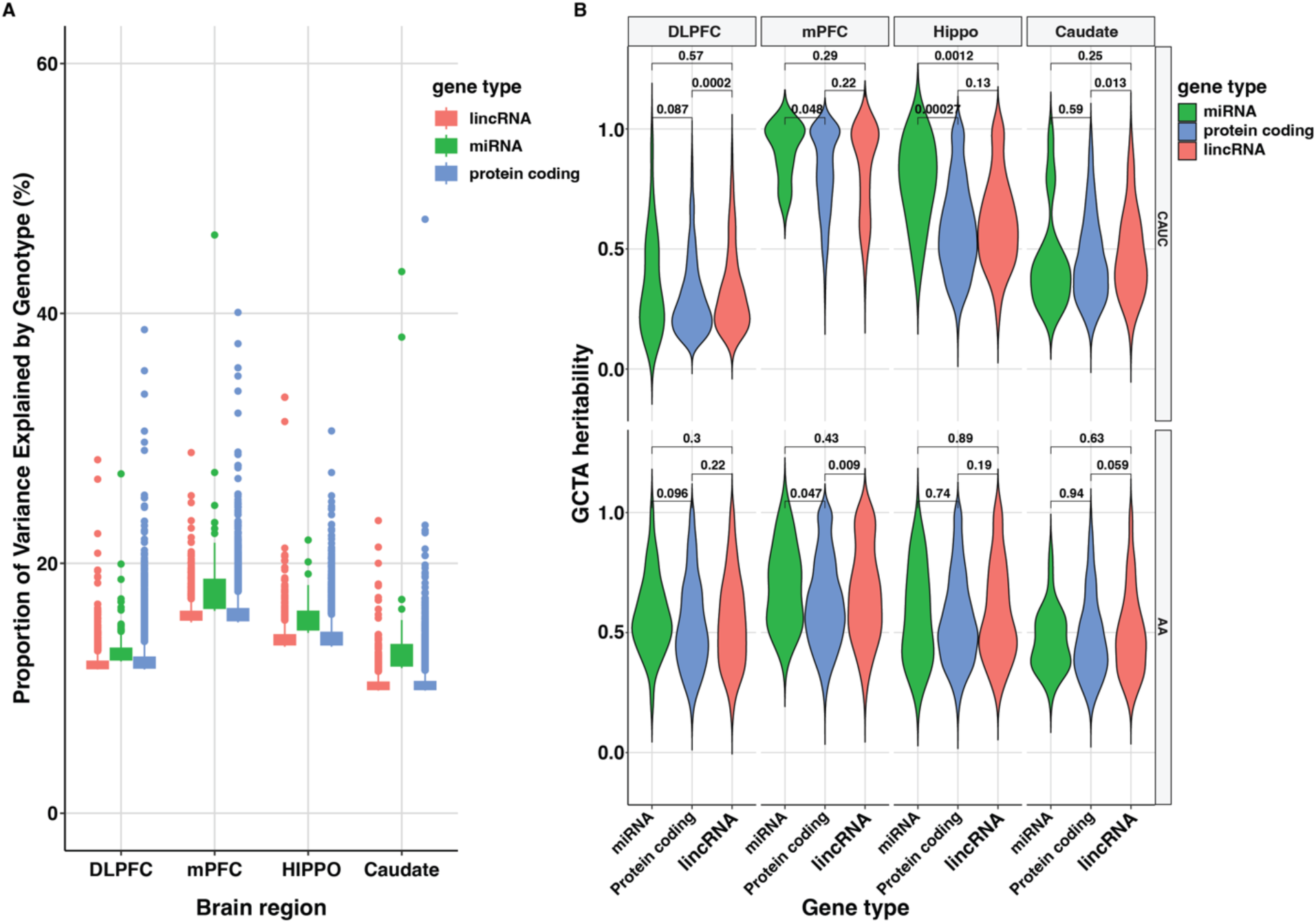
Distributions of miR-eQTL significant variant-miRNA pairs. (A) Distribution of the proportion of variance explained (PVE) by lead variants in African American (AA) ancestry. (B) Distribution of cis-heritability estimates of miRNAs calculated by GCTA.

**Supplementary Figure 8.**
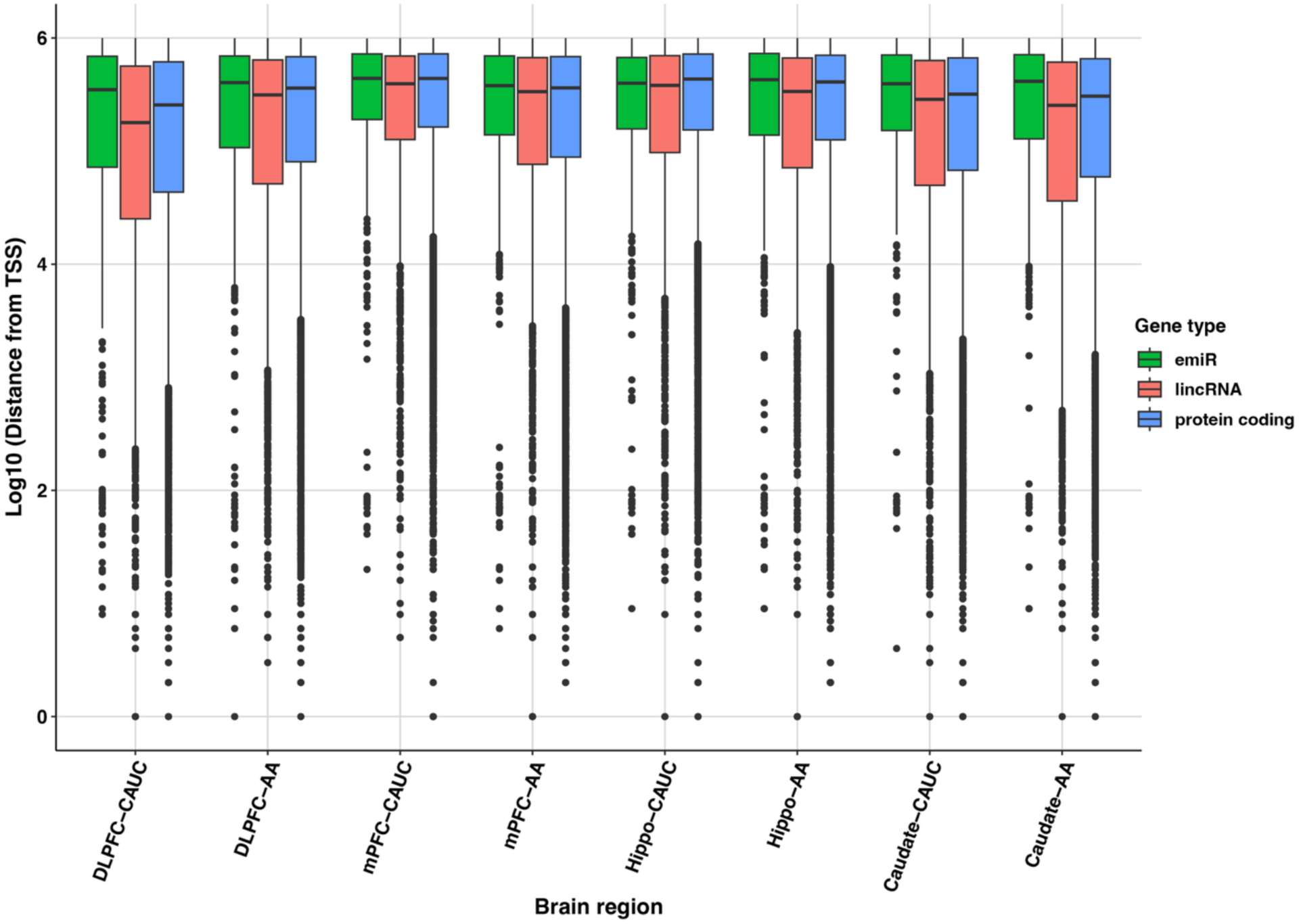
Distance between significant variants and transcription start sites (TSS). Distribution of distances between significant cis-eQTL variants and the TSS of their associated genes, including miRNAs, protein-coding genes, and lincRNAs. Distances are shown on a logarithmic scale.

**Supplementary Figure 9.**
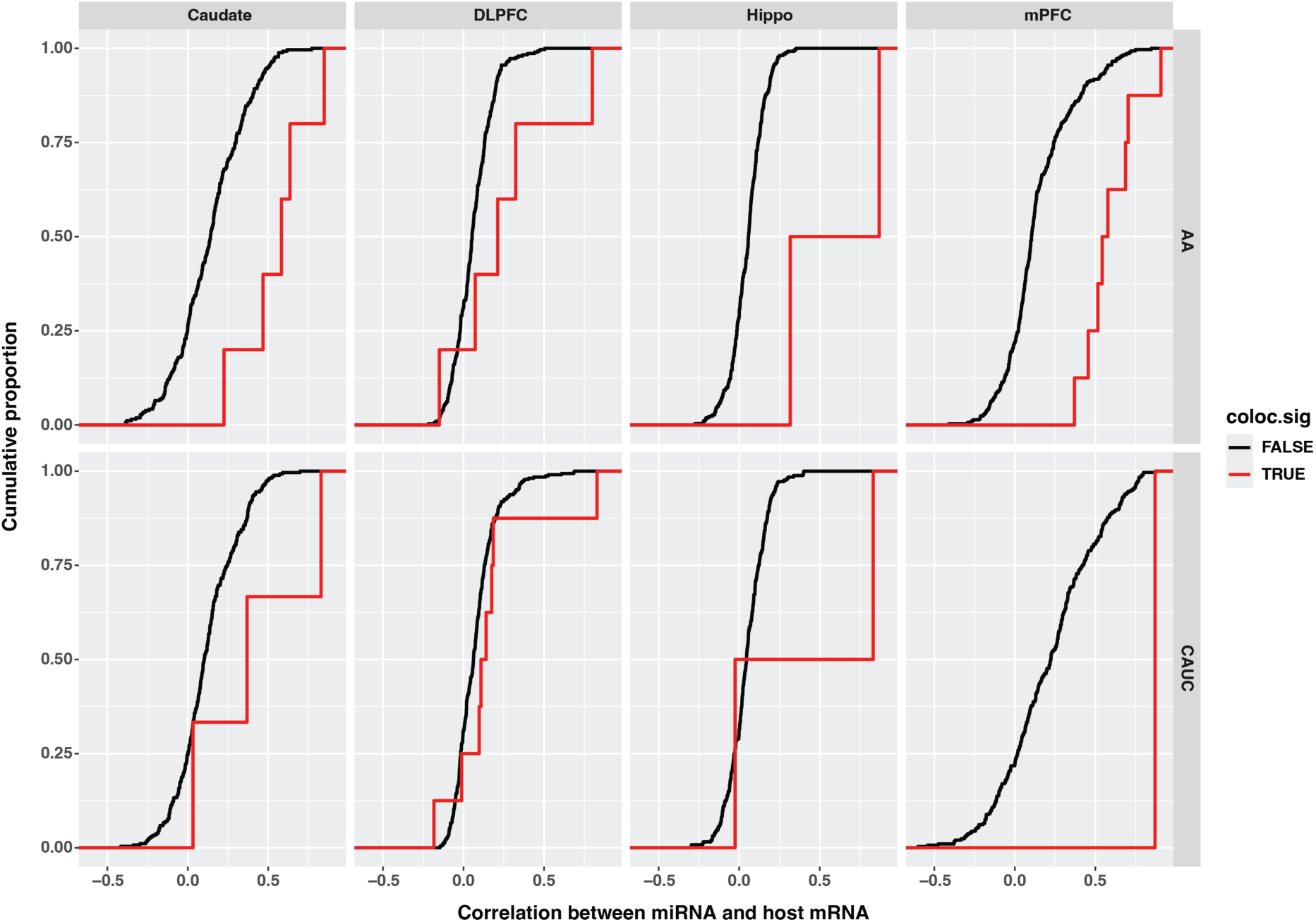
Correlation between intragenic miRNAs and their host genes. Distribution of Pearson correlation coefficients between intragenic miRNAs and their corresponding host protein-coding genes. Pairs showing significant colocalization based on coloc analysis (H₄ > 0.5) are highlighted in red.

**Supplementary Figure 10.**
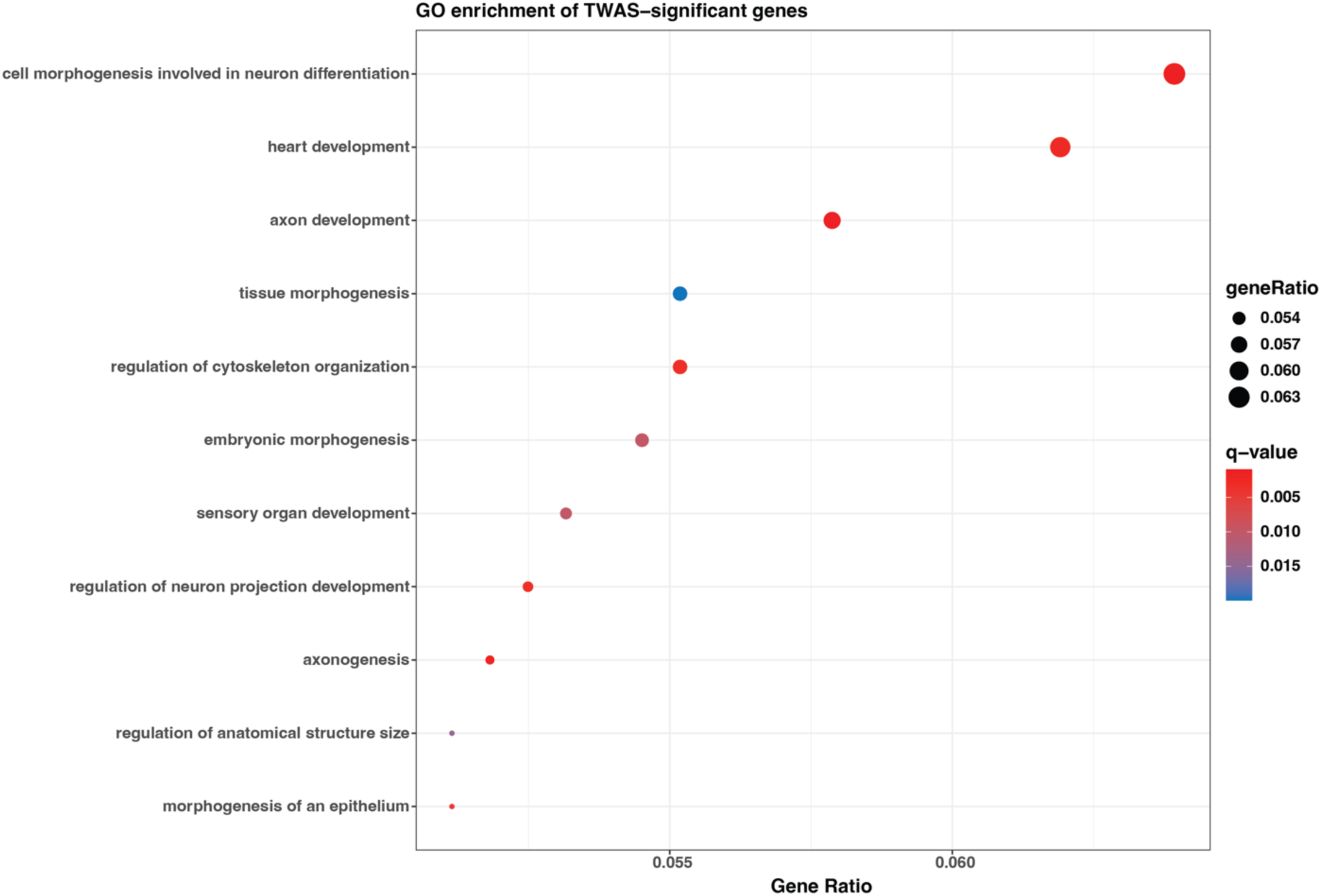
Significantly enriched Gene Ontology (GO) biological process terms for 18 miRNAs showing significant TWAS associations. The color gradient reflects –log₁₀(p-value).

**Supplementary Figure 11.**
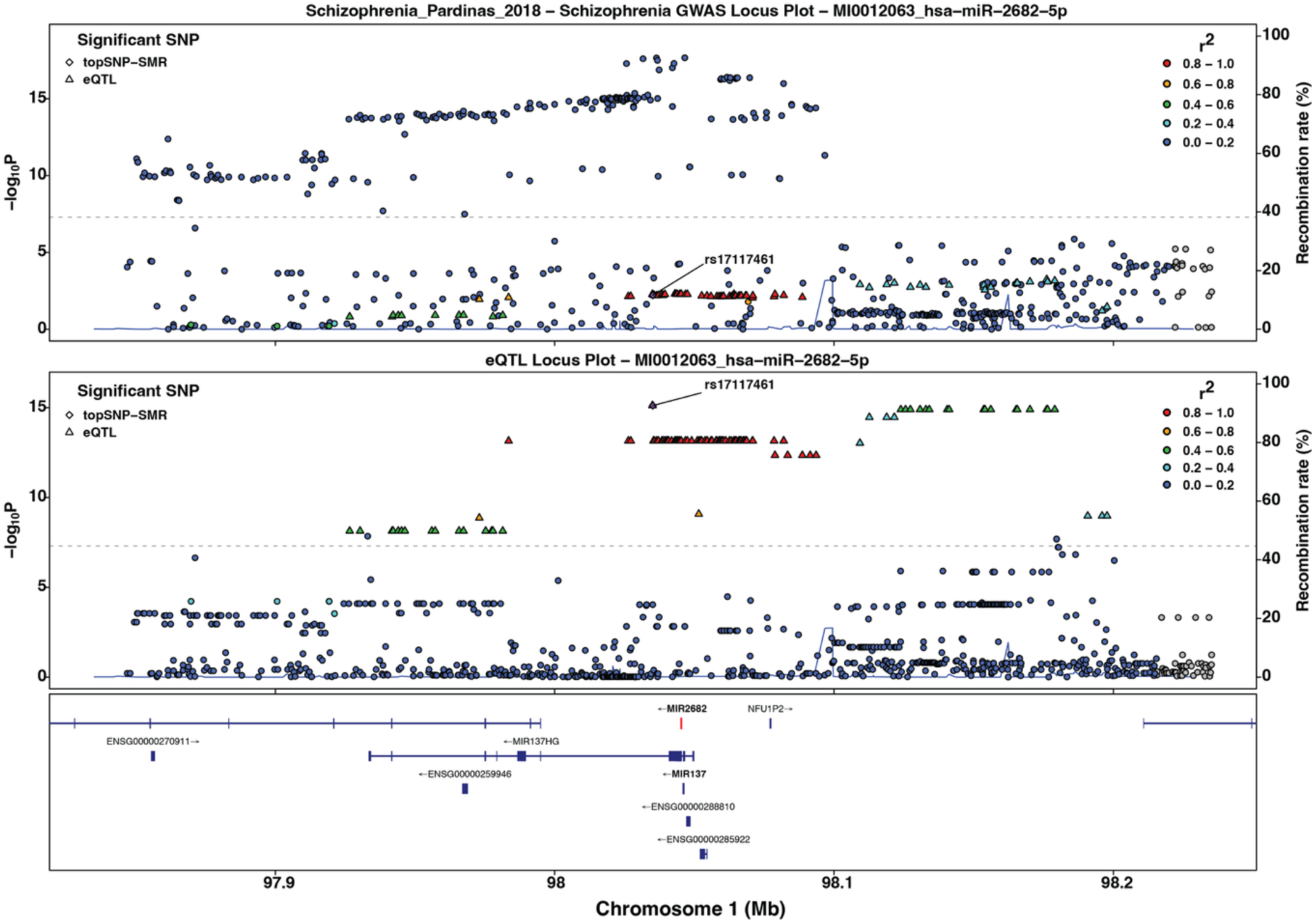
TWAS result for miR-2682-5p in the dorsolateral prefrontal cortex (DLPFC). The upper panel shows the corresponding GWAS association signals, and the lower panel shows the miR-eQTL signals.

**Supplementary Figure 12.**
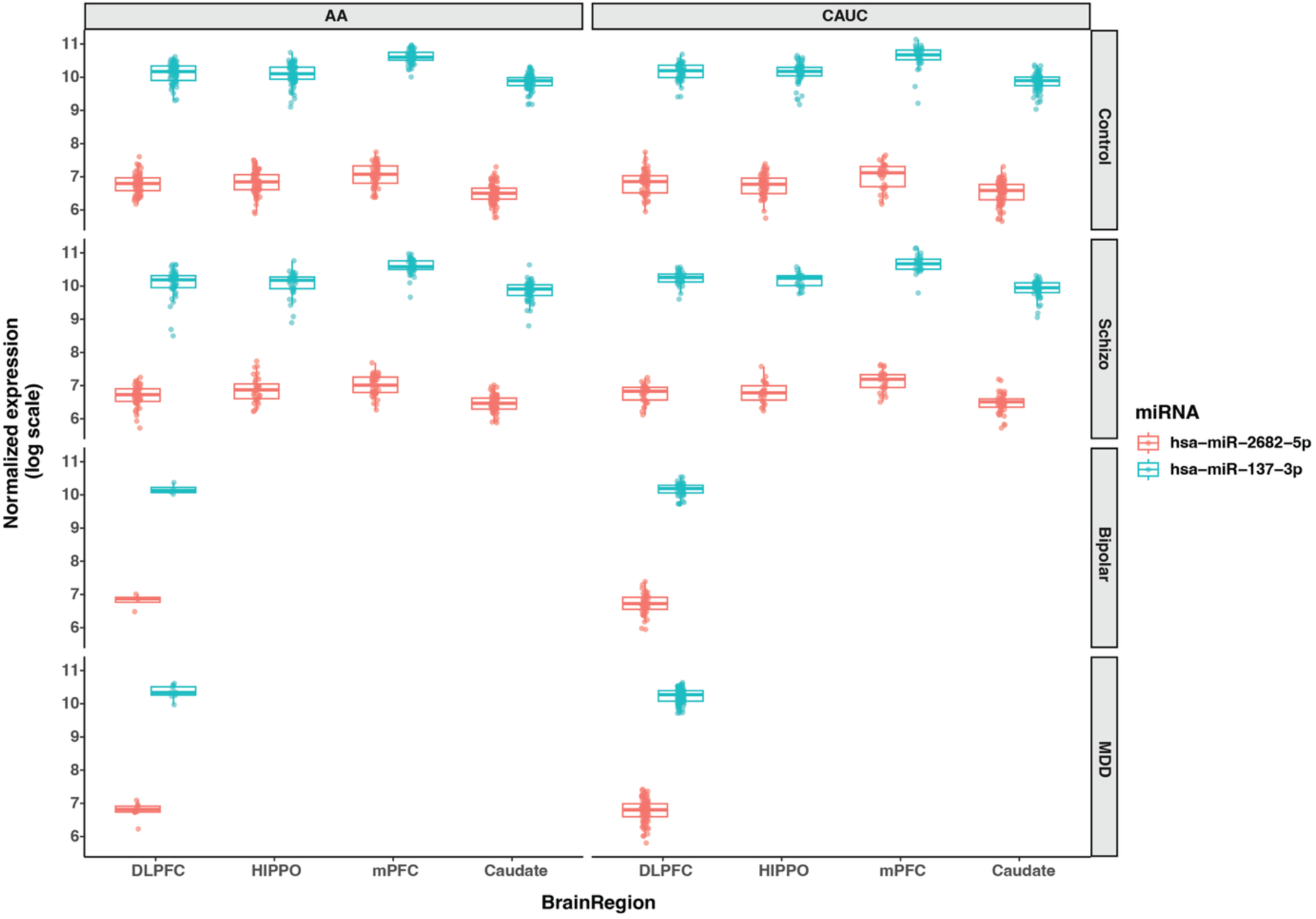
Expression levels of miR-137 and miR-2682. Depth-normalized read counts for miR-137 and miR-2682 across brain samples are shown.

